# The influence of visual pollution on navigation mechanisms in the damselfish (*Chrysiptera cyanea*)

**DOI:** 10.1101/2022.10.04.510829

**Authors:** William Michael Lunt, Theresa Burt de Perera, Cait Newport

## Abstract

Here, we investigate whether visual pollution has an effect on navigation in coral reef damselfish (*Chrysiptera cyanea*). Turbidity had no significant influence on the individual fish’s preference between egocentric and visual cues in a simple navigation task, with all individuals exhibiting a striking egocentric preference across all turbidity levels under testing. However, an alteration of cue preference may have occurred on a fine scale. Turbidity had profound effects on fish movement and decision-making behaviour, which has substantial implications for the behaviour of fishes on the ecological scale of a coral reef.

## INTRODUCTION

Coral reefs host a multitude of visually-guided behaviours. The reflection, absorption, polarisation and production of light are integral to behaviours underpinning the survival and reproduction of reef-dwelling organisms (Marshall et al. 2019; Losey et al. 1999; Kamermans & Hawryshyn 2011), from conspicuous territorial and breeding displays (Dawkins & Guilford 1994) to the use of landmarks for orientation (Braithwaite & Burt de Perera 2006). However, with rising coastal turbidity compromising the quality and spectral distribution of light (Jones et al. 2021), constricting photic zones (Cacciapaglia & van Woesik 2016) and restricting benthic irradiances (Anthony et al. 2004), visual behaviours are likely to be disrupted. In this study, we utilise a T-maze paradigm to interrogate the influence of turbidity on local navigation in a reef fish. Such an investigation is of the upmost importance if we are to understand how local navigation, a behaviour integral to the survival and fitness of reef fishes, is to be influenced in the coming decades of anthropogenic pollution.

Turbidity is a measure of the amount of light scattered by particles in a liquid upon a light being shone through that liquid (Kirk 1985). Acknowledged as an indicator of coastal water quality by the Australian and New Zealand Environmental and Conservation Council (ANZECC 2000), turbidity is rising in coastal ecosystems largely due to anthropogenic activities (Seers & Shears 2015). Owing to the intensification of meteorological and oceanographic processes (Cartwright et al. 2021), dredging (Collin & Hart 2015), algal blooms (Seers & Shears 2015), and the exacerbation of soil erosion and estuarine sediment discharge through the replacement of coastal forests with farmland (Wolanski & Spagnol 2000), coral reefs are subject to an increasing frequency and intensity of turbidity plumes.Extreme plumes have escalated to an intensity so as to be visible from space (Figure 1). Long-term analyses utilising both MODIS satellite images and in situ recordings reveal an upward trend in turbidity in tropical coastal waters when management strategies are absent (Seers & Shears 2015; Cartwright et al. 2021).

**Figure 1.**
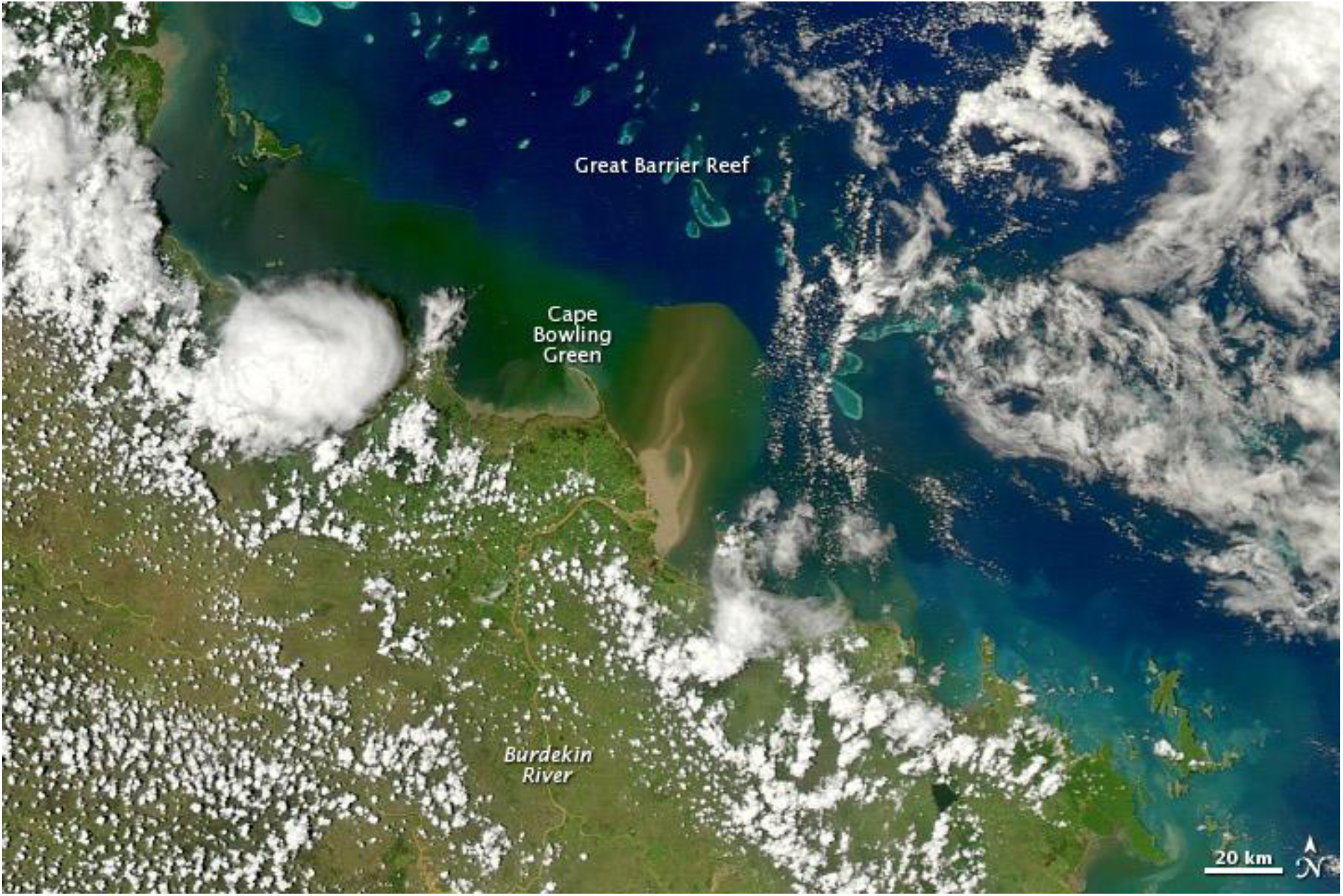
Satellite image showing a turbidity plume in Cape Bowling Green, Australia. This natural-colour image, captured by NASA’s Moderate Resolution Imaging Spectroradiometer (MODIS) on 4 January 2011, shows an extreme turbidity plume in Cape Bowling Green viewed from space, generated by sediment discharge from the Burdekin River. Image courtesy of Norman Kuring. Annotations courtesy of Michon Scott (NASA Goddard Space Flight Center 2014).

Turbidity has profound effects on light in aquatic systems (Bowers & Binding 2006). The materials which contribute to turbidity are categorised into chromophoric dissolved organic matter (CDOM), phytoplankton and non-algal particulates (NAP) (Jones et al. 2021). While water absorbs strongly in the longer red wavelengths (Kirk 2010), phytoplankton absorbs strongly in the shorter blue-green wavelengths (Van Duin et al. 2001), CDOM in the shorter blue-ultraviolet wavelengths (Shi et al. 2013) and NAP in the shorter blue wavelengths with a tapering absorbance towards the longer red wavelengths (Babin et al. 2003). As a consequence of these different materials differentially truncating the distribution of wavelengths reaching reef communities, one would expect different light-dependent behaviours and processes to be impacted differently by different sources of turbidity (Nieman et al. 2018). In addition to altering the spectral distribution of light, turbidity constricts irradiance, accounting for up to 80% of total annual variation in benthic irradiance and primarily governing temporal cycles of irradiance in coral reefs (Anthony et al. 2004).The effect of such shading depends on its extent. Moderate levels of turbidity can be expected to confer shaded refuge to corals otherwise exposed to the harmful interaction between irradiance and sea surface temperature, while persistent, extreme turbidities may preclude up to 16% of otherwise viable coral habitat through shading (Cacciapaglia & van Woesik 2016).

A growing body of literature is revealing the consequences of turbidity for non-visual and visual behaviours and processes alike in teleosts. Among the non-visual influences of elevated turbidity are heightened plasma cortisol (Au et al. 2004), avoidance behaviour (Collin & Hart 2015), reduced movement efficiency (Newport et al. 2021), gill damage (Wong et al. 2013; Au et al. 2004), non-linear changes in predator-prey interactions (Wenger et al. 2013), reduced foraging, growth and activity (Wenger et al. 2012; Leahy et al. 2011), and modifications of social structure (Borner et al. 2015). Beyond teleosts, waterbird diving behaviour (Thompson & Price 2006) and distribution (Henkel 2006), harbour seal visual acuity (Weiffen et al. 2006) and copepod community structure (Liljendahl-Nurminen et al. 2008) are also influenced. Among visual behaviours, turbidity was found to reduce reaction distances (Vogel & Beauchamp 1999; Sweka & Hartman 2003; Collin & Hart 2015; Meager et al. 2005), disrupt visually-guided settlement behaviour of larvae (Wenger et al. 2011), induce a plastic shift in the expressed opsin repertoire to favour motion detection (Ehlman et al. 2015), and to increase reliance on non-visual senses (Suriyampola et al. 2018; Meager et al. 2005). To expand on this final point, this sensory compensation, whereby an animal shifts their primary sensory modality in response to the quality or reliability of information of different modalities, has been documented across a broad range of taxa (Heuschele et al. 2009; Ryugo et al. 1975; Lessard et al. 1998; Suriyampola et al. 2018) and likely has widespread ethological implications in coral reefs.

Navigation is one such behaviour likely to be perturbed by elevated turbidity. Fish concurrently utilise a range of cues to orient and navigate in their local environment, including beacons, whereby a landmark directly marks the location of a goal (Warburton 1990), ordered lists of landmarks, whereby the fish follows a chain of ordered landmarks in a process called pilotage or chaining (Burt de Perera 2004), and body-centred, egocentric cues, whereby the fish utilises vectors and movement parameters relative to their own body to compute distance and direction (Rodríguez et al. 1994; Odling-Smee & Braithwaite 2003a; Braithwaite & de Perera 2006). Given there is usually no conflict in the information provided by different cues, multiple cues may be used jointly when making navigational decisions (Braithwaite & de Perera 2006). Such multi-cue integration was neatly demonstrated by Sovrano and colleagues (Sovrano et al. 2003) through the combined use of geometric and visual cues by redtail goodeids *Xenotoca eiseni* (Rutter 1896) to locate a goal. Similarly, fish have been found to independently use and integrate egocentric and landmark-based cues when faced with navigational tasks (Burt de Perera & Garcia 2003; Odling-Smee & Braithwaite 2003b).

While fish have the capacity to utilise multiple cues of different modalities independently or concurrently for navigation, that is not to say cues of different modalities are used equally. Sutherland and colleagues (Sutherland et al. 2009) found that sighted cavefish *Astyanax fasciatus* (Cuvier 1819) exhibited a preference for following visual landmarks when landmark cues were placed in conflict with egocentric turning cues, despite each modality having provided equally reliable information during training. Such modal preferences likely stem from a combination of the organism’s evolutionary history and their life history (Odling-Smee & Braithwaite 2003b; Bensky & Bell 2018). In support of this, when faced with a T-maze task where egocentric cues were in conflict with landmark cues, three-spined sticklebacks *Gasterosteus aculeatus* L. sourced from ponds relied more on visual landmarks than sticklebacks from rivers did (Odling-Smee & Braithwaite 2003b). This heightened preference for visual cues in pond-dwellers is likely due to landmarks providing fixed, reliable locational information in ponds, while in rivers it is less likely that any salient landmark will remain in the same place for long (Odling-Smee & Braithwaite 2003b).

With turbidity imparting such substantial changes to the spectral environment in coral reefs, one would expect a visually-guided behaviour such as navigation to be influenced. Yet, despite navigation underpinning each of foraging, reproduction and survival for many reef fishes, little work has explored the influence of turbidity on navigation in reef fishes. Newport and colleagues (Newport et al. 2021) explored the influence of elevated turbidity on the movement behaviour of Picasso triggerfish *Rhinecanthus aculeatus* L. in a foraging task, finding individuals took slower, less efficient foraging routes in high turbidity. Such reductions in movement efficiency are likely to be detrimental to the survival and consequent population dynamics of triggerfish and other visually-guided reef fishes alike (Newport et al. 2021).Similarly, in a freshwater context, Sekhar and colleagues (Sekhar et al. 2019) trained fish to find food pellets in the presence or absence of landmarks, in a range of turbidity conditions. While feeding latencies did increase with turbidity, this increase was significantly smaller when visual landmarks were present, indicating the zebrafish were utilising the brightly coloured landmarks for guidance in turbid conditions (Sekhar et al. 2019).

In this study, I sought to elucidate i) the influence of turbidity on the preference of reef fishes between different navigational cues, and ii) to explore how fish movement is affected by elevated turbidity. With regards to the first objective, one would expect that as turbidity rises and the quality and accessibility of visual information degrades, fish would come to rely on body-centred, egocentric cues over visual cues for navigational guidance. However, such a shift in cue preference is likely to be non-linear, with an increased attentiveness for visual landmarks in low turbidities for guidance, as in Sekhar et al. (2019), but a reduced attentiveness for visual landmarks in high turbidities when they are no longer visible. With regards to the second objective, with elevated turbidity limiting visual information on navigational cues and potential danger cues, one would expect fish to take slower, more hesitant, tortuous paths in search of food. To address these objectives, six damselfish *Chrysiptera cyanea* (Quoy & Gaimard 1825) were trained in a T-maze to turn in a given direction in the presence of a visual landmark. In probe trials, egocentric cues and landmark cues were placed in conflict under a range of turbidities to explore the influence of turbidity on the cue preference of reef fishes in this navigational task. Movement trajectories were produced for each fish for every trial, allowing an examination of the influence of turbidity on their movement. Such a study is of the upmost importance if we are to understand whether turbidity is inducing shifts in navigational cue preference in reef fishes, and whether such shifts are sufficient to maintain their efficiency in navigational tasks.

## MATERIALS & METHODS

### SUBJECTS AND HUSBANDRY

Six blue devil damselfish (*Chrysiptera cyanea*) were used in this experiment. This sample size provides adequate power under a generalised linear mixed effects model (GLMM) utilising a binomial distribution (Supplementary Information A). All fish were experimentally naïve, adult females caught from the Great Barrier Reef and purchased from a local supplier (The Goldfish Bowl, Oxford). Each fish was housed in an individual aquarium (35×30×60cm, 19cm water depth, 39.9L filled volume) equipped with a filter pump (EHEIM biopower 160 or EHEIM aqua 160), internal aquarium heater (V2therm 50 Digital Heater 50W), and rocks, gravel and a plant for habitat enrichment. Fish were visually isolated from each other to prevent behavioural interactions. Fish were fed daily on a diet of crisps (TetraMin, Crisps Fish Food) and mysis shrimp (Gamma blister, Mysis Shrimp). Water parameters were tested regularly and water changes carried out accordingly. Tank water was treated with Vetark Fluke-Solve (0.0038g/L) to kill parasites. A log of fish appetite and activity was completed bidaily to monitor fish health and condition.

### APPARATUS

An opaque plastic T-maze was placed into the home tank of each individual during pre-training, training and experimental sessions. A guillotine door was secured at the T-maze entrance to control when the fish could access the maze (Figure 2). The region which the fish resided in between trials was termed the inter-trial area. A grey LEGO tower (80×16×16mm) was used as the visual landmark in pre-training, training and experimental sessions, with a plastic sheet at its base to allow it to be hooked under the edge of the T-maze, securing it in place. A video camera (GoPro Hero 8, FPS: 60; Resolution: 1080p; FOV: linear) was mounted 37.5cm above the water surface. Tanks were lit using i) strips of blue-white LEDs positioned at the back of each tank and, during filming, ii) two softbox lights (Neewer700W 24”x24” Softbox with 85W 5500K CFL Light Bulb). No lights were positioned directly above tanks to avoid reflections on the water during recordings.

**Figure 2.**
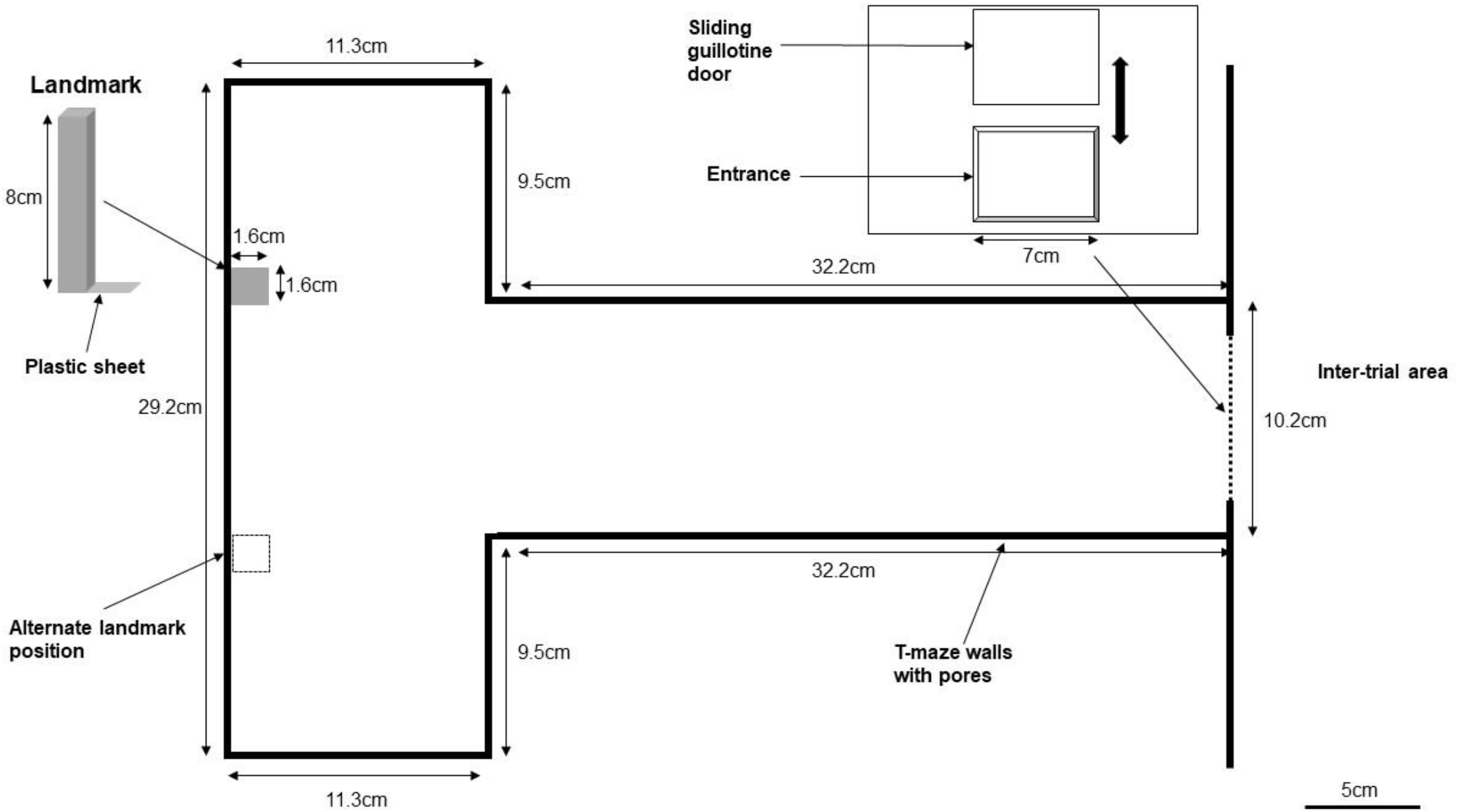
T-maze schematic. This schematic shows the apparatus used in the pre-training, training and experimental phases of this study. Before a session, the T-maze was placed into the tank of an individual with the arms of the T-maze positioned at the opposite end of the tank to the inter-trial area. The walls of the T-maze were perforated with small holes to allow turbidity to evenly distribute across the tank. A plastic sheet housing the sliding guillotine door, which was operated using a pulley system, was placed at the base of the stem of the T-maze and secured in place using clips. The width of this plastic sheet was equal to that of the tank, meaning fish were restricted to the inter-trial area once this sheet was positioned in the tank before the guillotine door was opened. The LEGO landmark could be placed in one of two positions in the maze, as shown by the displayed position of the landmark and by the alternate landmark position.

### TRAINING

Individuals were trained to navigate a T-maze to receive a food reward. Five individuals were randomly assigned to be trained to turn left (FishID=1;2;4;9;10), and five individuals right (FishID =3,5,6,7,8). Two individuals learnt to turn left (FishID=4;9), and four individuals right (FishID=3;5;6;7). For individuals trained to turn left, the landmark was placed on the back wall of the T-maze at the beginning of the left arm, and the individual was rewarded upon their caudal fin passing into the left arm. The opposite applied for individuals trained to turn right. Individuals were trained to an 80% performance threshold, meaning they were deemed to have learnt the task when, for three consecutive training sessions (whereby a training session entails 10 trials), the individual took the correct arm in at least 80% of trials. A trial was terminated if the individual had not entered the maze after 300 seconds, or when they had taken a turn and been rewarded (if they took the correct turn) or not rewarded (if they took the incorrect turn) and the guillotine door had been shut. The duration (s) of a training trial constituted the time for which the door was open. Approximately one minute was left between consecutive training trials. Trials were filmed. Two pre-training phases were used to accelerate training (Supplementary Information B).

### EXPERIMENTAL TREATMENTS

Five turbidity treatments were used. These conditions encompass the range of turbidities that Indo-Pacific damselfish encounter in the wild (Wolanski et al. 2008; Orpin & Ridd 2012). Bentonite clay was added to each tank to achieve the desired turbidity, chosen for its presence in coastal environments and ease of suspension. The treatments were i) Turb0 (Control) (0g/L), ii) Turb1 (0.063g/L), iii) Turb2 (0.125g/L), iv) Turb3 (0.201g/L), and v) Turb0Sys (0g/L). Turb0Sys serves to detect potential influences of the systematic increase in turbidity employed in this experiment, as detailed under ‘Experimental Procedure’.Turbidities were measured using a turbidimeter (HACH2100Q Portable Turbidimeter). The range of turbidities (nephelometric turbidity units; NTU) generated at each turbidity level may be found in the Supplementary Information (Supplementary Information C).

### EXPERIMENTAL PROCEDURE

During each experimental session, six trials were conducted. This entailed four unrewarded experimental trials, followed by two training trials in which a food reward was provided. No more than four experimental trials were conducted in a row to avoid extinction of the learnt behaviour or a loss of motivation. The two training trials served to reinforce the learnt behaviour after that day’s testing. Training trials did not precede experimental trials in order to prevent the following of odour trails in experimental trials.

During each experimental trial, the landmark was positioned in one of two configurations: N (normal) or Opp (opposite). In the N configuration, the landmark was positioned in the arm which that fish had been trained to take. In the Opp configuration, the landmark was positioned in the opposite arm to that which the fish had been trained to take. Consequently, the information provided by the landmark and the information provided by the egocentric turning cue were in conflict when the landmark was in the Opp configuration. Therefore, which cue the individual follows (landmark or egocentric) in a trial with the Opp configuration elucidates which cue they prioritise under those conditions. Both N trials and Opp trials are needed to elucidate the influence of landmark position on turning behaviour. During each session, there was balance in landmark configuration (two N trials and two Opp trials per session).

For an experimental trial, the T-maze was placed into the individual’s tank, with the individual residing in the inter-trial area. To start the experiment, the guillotine door was lifted and the fish allowed to navigate the maze. Upon an individual completing a trial, defined as that individual’s caudal fin passing into an arm of the maze, the experimenter waited a short time before (if required) gently ushering the fish back into the inter-trial area, before closing the guillotine door. The next trial began 1 minute after the fish returned to the inter-trial area. The latency to enter the maze, latency to take an arm and turn direction were recorded.

Experimental trials were conducted under each of the five turbidity treatments. A known mass (g) of bentonite clay was added to each tank at the beginning of the testing period for that turbidity level, and gently mixed by hand prior to each session. Turbidity was recorded before and after the experimental session for that fish. In accordance with the power analysis, sixteen experimental trials were conducted per fish per treatment (n=480).

While randomisation of treatment order within each experimental session would be preferable, this was logistically impractical given the constraints of the system used. All testing and consequent manipulation of turbidity had to occur in an individual’s home tank as damselfish experience high stress following tank transfers. This prevented the utilisation of dedicated experimental tanks within which one could generate each turbidity level independently and move fish between for each trial. Turbidity could not be decreased from one trial to the next as this required a complete (39.9L) water change which rapidly depleted the 200L saltwater reservoir available, which takes multiple hours to re-stock, preventing testing of all fish on the same day. Running an internal filter to remove clay particles between trials was also insufficient. Given these constraints, it was concluded that experimental trials would be conducted in a systematic manner, with all Turb0 tests being conducted first, followed by all Turb1 tests, followed by all Turb2 tests, followed by all Turb3 tests. Upon finishing testing for each turbidity level, tanks received a quarter water change to maintain optimal water conditions. Upon finishing Turb3 testing, tanks received a half water change and tank clean, restoring water to 0NTU. To account for potential influences of this systematic increase in turbidity, fish were tested at 0NTU again following the completion of the Turb3 tests. This final treatment was termed Turb0Sys. Within-individual comparisons of the two sets of results obtained under 0NTU conditions, namely from before (Turb0) and after (Turb0Sys) exposure to the systematic turbidity increase, would allow detection of behavioural changes that may have occurred in response to the systematic turbidity increase.

### MOVEMENT TRACKING

ToxTrac, an open source tracking software with a history of application to fish, was used to track and analyse the trajectories of fish during each trial (Rodriguez et al. 2018). Videos were converted to .avi format with 30 FPS and 720p resolution for use in ToxTrac. ToxTrac was calibrated for each fish independently by inputting six images of a 20×14 grid of black and white squares placed on the bottom of the tank and specifying the width (mm) of each square. Calibration was necessary for the software to translate pixels into millimetres. To assess the accuracy of ToxTrac’s inbuilt calibration feature, a manual calibration was conducted. A 2% discrepancy was detected between ToxTrac’s automatically calculated pixel:mm conversion rate and the manually calculated pixel:mm conversion rate. This discrepancy was deemed satisfactorily small given the short distances that fish were operating over.

For each video (n=480), the experimenter manually adjusted tracking parameters (Object Colour Selection, Min/Max Object Size, Max Displacement/Frame) to optimise detection of the fish. Six movement parameters were extracted from each analysis: a) average speed (mm/s), b) average mobile speed (mm/s), c) average acceleration (mm/s^2^), d) total distance travelled (mm), e) exploration rate and f) mobility rate. Grid-use frequencies were also extracted to elucidate how the arena space was being used.

### STATISTICAL ANALYSIS

#### Navigational cues

To elucidate the influence of turbidity on the preference between egocentric and landmark cues, a GLMM utilising a binomial distribution was constructed (‘lme4’ R package). The ‘generalised’ aspect of a GLMM was required to model the binary data of 100s (whereby a score of 100 for a trial indicated the fish turned in the egocentric direction) and 0s (whereby a score of 0 for a trial indicated the fish turned in the landmark direction). The ‘mixed’ aspect was required to account for repeated measures within each fish. Turbidity level was modelled as a fixed effect. FishID and the post-testing turbidity were modelled as random effects to account for, respectively, repeated measures within each fish and for any fall in turbidity that occurred over a session. This analysis was conducted utilising data exclusively when cue conflict was present (Opp trials).

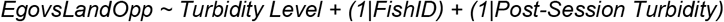

A second GLMM was constructed to elucidate the influence of landmark configuration on the preference between egocentric and landmark cues, and any potential interaction landmark configuration may have with turbidity. This model partitioned data regarding which turn the fish took depending on landmark configuration and compared the difference in percentage of egocentric turns between N trials and Opp trials with a ‘no difference’ value of 0.

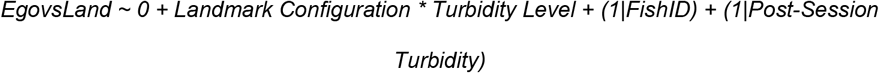

#### Trajectory analysis

To detect differences in the six movement parameters between turbidity levels, linear mixed effects models (LMMs) were constructed. Landmark configuration (and a potential interaction with turbidity) was incorporated to explore whether the landmark was attended to. The same analysis was utilised for decision hesitance: the time (s) between entering the maze and taking a turn.

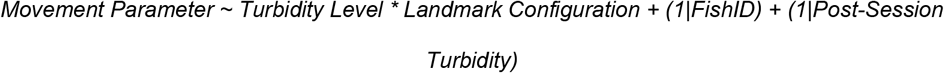

To explore how the maze area was used during experimental trials in response to landmark configuration, an LMM was constructed. This model explored the influence of landmark configuration on the proportion of time spent in each XY bicoordinate in the T-maze.

Turbidity level was modelled as a random effect to account for any influence turbidity may have on the attendance of fish to the landmark cue.

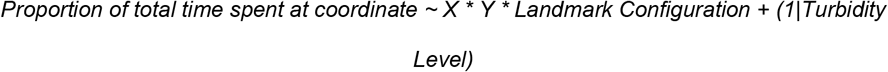

#### Speed-efficiency trade-off

Within each navigational decision is a trade-off between the speed and efficiency of that decision. To elucidate the influence of turbidity on the range of solutions adopted in speed-efficiency trade-off space, the decision time (decision speed) was plotted against the total distance travelled (decision efficiency) for each trial for each fish at all turbidity levels. A 95% confidence interval ellipse was drawn to encompass the points in trade-off space for each turbidity level (Figure 3). Outliers were specified (*Outlier< Q1– 1*.*5xIQR, Outlier>Q3+1*.*5xIQR)* and removed prior to the drawing of ellipses. A linear model analysed the influence of turbidity on the area of these ellipses, whereby the area of an ellipse quantified the range of solutions adopted in speed-efficiency trade-off space at that turbidity level. The assumptions of a linear model were met following a cube transformation of ellipse areas.

**Figure 3.**
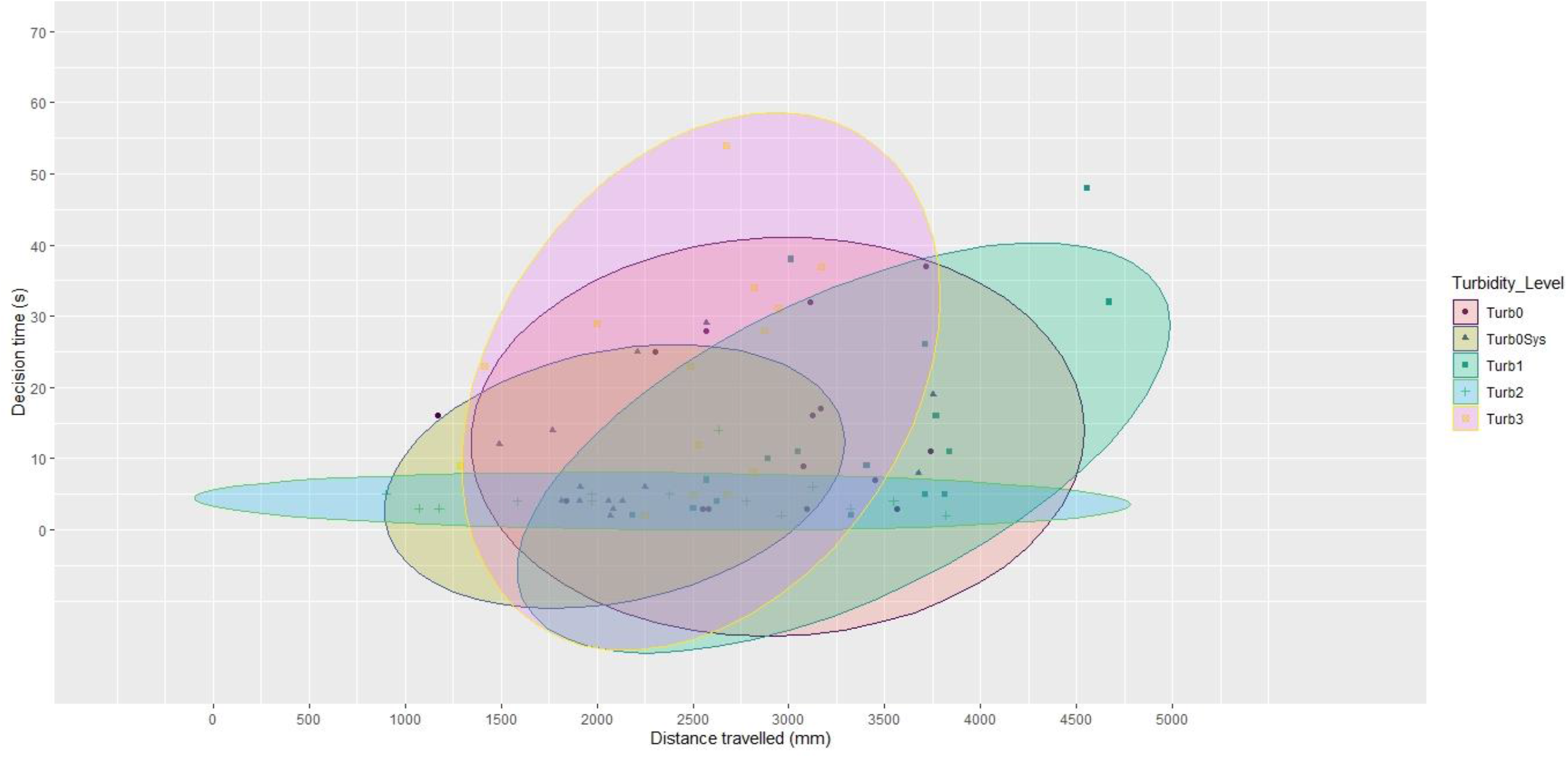
Plot of distance travelled (mm) against decision time (s) for every trial for Fish 3, with 95% confidence interval ellipses drawn at each turbidity level. Data from *Chrysiptera* cyanea. This plot shows the total distance travelled (mm) plotted against the decision hesitance (s) for each trial across the five turbidity levels for Fish 3. Similar graphs may be seen for each of the five other fish (Supplementary Information D-H). Each point corresponds to a distinct trial for that fish. The shape of the point indicates the turbidity level to which that trial corresponds. A 95% confidence interval ellipse was drawn for each turbidity level to encompass the spread of points at that turbidity level. Consequently, five ellipses are drawn per fish. Outliers were removed prior to the production of this graph.

To elucidate the influence of turbidity on the distribution of solutions in speed-efficiency trade-off space, a linear model analysed the effect of turbidity on the overlap area between ellipses. Should the area of overlap between two ellipses be significantly lower than that between two ellipses in clear water, the distribution of solutions in trade-off space must be shifting with a change in turbidity (Figure 4). For example, should one find that the overlap area between a Turb0 ellipse and a Turb3 ellipse is significantly smaller than that between a Turb0 ellipse and a Turb1 ellipse, this suggests an increase in turbidity is causing the spread of solutions in trade-off space to move away from that found in clear water. Overlap areas were calculated for each pairwise turbidity level comparison. Following a square-root transformation of overlap areas, the assumptions of a linear model were met.

**Figure 4.**
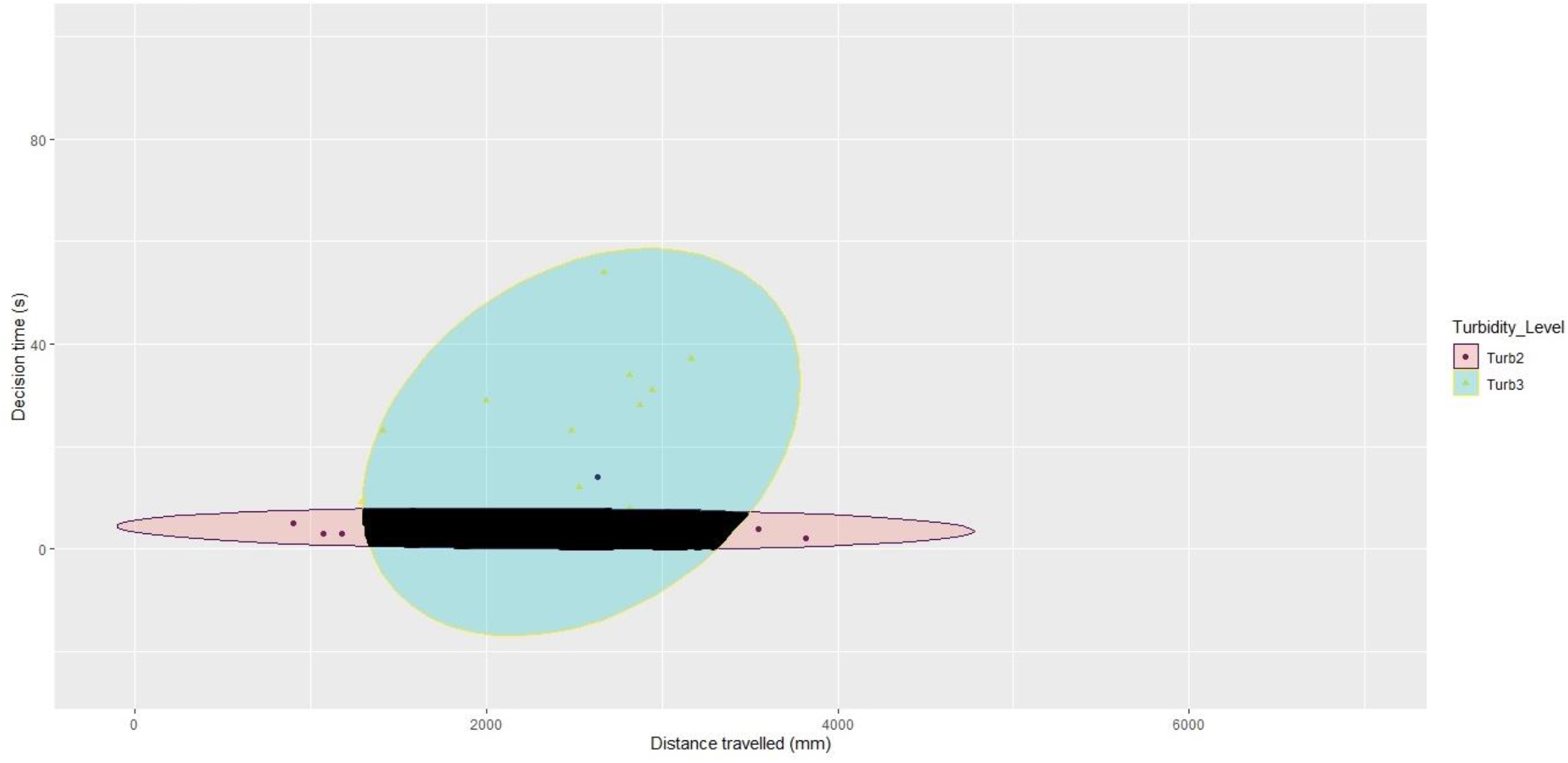
Plot of distance travelled (mm) against decision time (s) for trials in Turb2 and Turb3 for Fish 4, with 95% confidence interval ellipses drawn for each turbidity level and the area of ellipse overlap highlighted in black. Data from *Chrysiptera* cyanea. This plot shows the total distance travelled (mm) plotted against the decision hesitance (s) for each trial in Turb2 and Turb3 for Fish 4. This combination of turbidities and fish ID was randomly selected as an example. Each point corresponds to a distinct trial for that fish. The shape of the point indicates the turbidity level to which that trial corresponds. A 95% confidence interval ellipse was drawn for each turbidity level to encompass the spread of points at that turbidity level. The black region highlights the area of overlap between these ellipses. It is this overlap area which is utilised for the overlap area analysis outlined under the ‘Speed-efficiency trade-off’ subheading.

### ETHICAL CONSIDERATIONS

The care and use of experimental animals complied with the animal welfare laws, guidelines and policies of The University of Oxford AWERB (Permit Reference Number: APA/1/5/ZOO/NASPA/Newport/Damselfish). To minimise stress associated with changes in turbidity, the range of turbidities used was kept within that which fish experience in the wild.

## RESULTS

### Navigational cues

The results of the six trained fish indicate that turbidity does not cause a shift in cue preference. A GLMM reported no significant effect of turbidity on the percentage of turns in the egocentric direction when egocentric and landmark cues were in conflict, with no significant difference in the percentage of turns in the egocentric direction between Turb0 and Turb0Sys (*z* = 0.390, *p* = 0.695), Turb1 (*z* = 0.304, *p* = 0.760), Turb2 (*z* = 0, *p* = 1.000) or Turb3 (*z* = 1.234, *p* = 0.215) (Figure 5). All fish exhibited similar variability in the percentage of egocentric turns taken (*F*_*1,4*_ = 0.6257, p=0.473) (Supplementary Information I), and none exhibited innate directional biases (Supplementary Information J).

**Figure 5.**
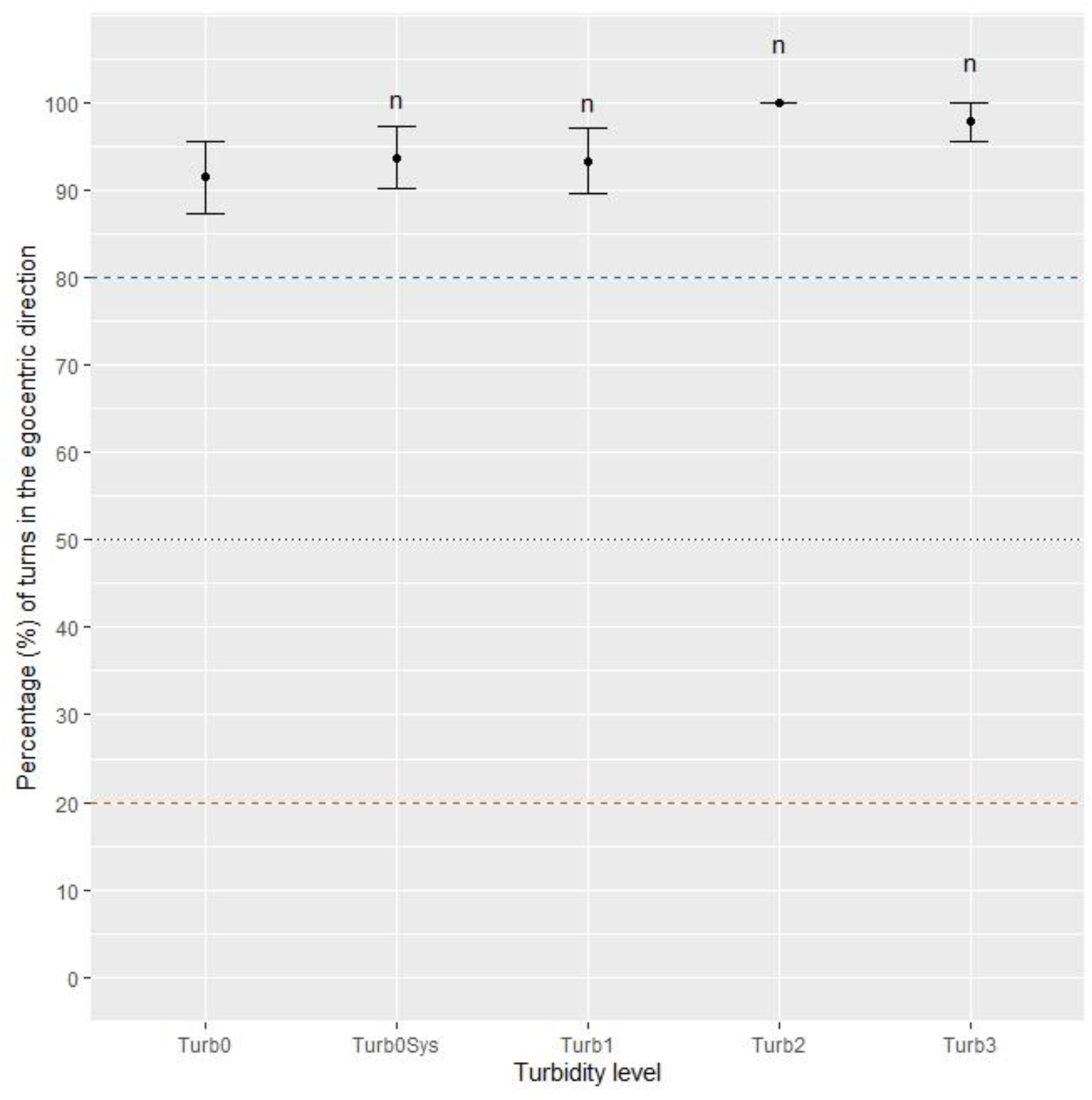
Plot of mean percentage (%) of total trials where cue conflict was present in which fish followed their learnt egocentric cue, at each turbidity level. Data from *Chrysiptera* cyanea. The plot shows the percentage (%) of the total trials where cue conflict was present in which the fish took the arm corresponding to their learnt egocentric cue. Errors bars indicate 95% confidence intervals about the mean. The letter n above a point indicates that that value is not significantly different from the corresponding value for Turb0. A value of 100% indicates fish followed egocentric cues in every trial at that turbidity level. A value of 0% indicates fish followed landmark cues in every trial at that turbidity level. The black dotted line with intercept 50% represents where we would expect points to lie were the fish behaving randomly or weighing egocentric and landmark information equally. The blue dashed line with intercept 80% represents where we would expect points to lie were fish completely prioritising egocentric cues, given the 80% performance criterium in training. The brown dashed line with intercept 20% represents where we would expect points to lie were fish completely prioritising landmark cues, given the 80% performance criterium in training.

Landmark configuration had no significant effect on preference between egocentric cues and landmark cues at any turbidity level (*z* = -0.803, *p* = 0.422). In support of this, when the landmark was in the Opp configuration, there was no significant difference in the percentage of egocentric turns between Turb0 and Turb0Sys (*z* = 0.394, *p* = 0.695), Turb1 (*z* = 0.745, *p* = 0.760), Turb2 (*z* = 0.000, *p* = 1.000) or Turb3 (*z* = 1.259, *p* = 0.215). Similarly, no significant difference in the percentage of egocentric turns was reported between Turb0 and Turb0Sys (*z* = 0.576, *p* = 0.568), Turb1 (*z* = -0.828, *p* = 0.408), Turb2 (*z* = 0.000, *p* =1.000) or Turb3 (*z* = -0.828, *p* =00200.410) when the landmark was in the N configuration (Figure 6). Despite this, the data do show the hypothesised pattern entailing a negative effect of the Opp configuration in clear water, whereby a negative effect constitutes the landmark arm being taken more often in Opp trials than in N trials at that turbidity, and a negligible or positive effect of the Opp configuration at higher turbidities where the landmark cannot be seen clearly (Figure 7). In summary, visual pollution had no significant effect on which cues were attended to during local navigation.

**Figure 6.**
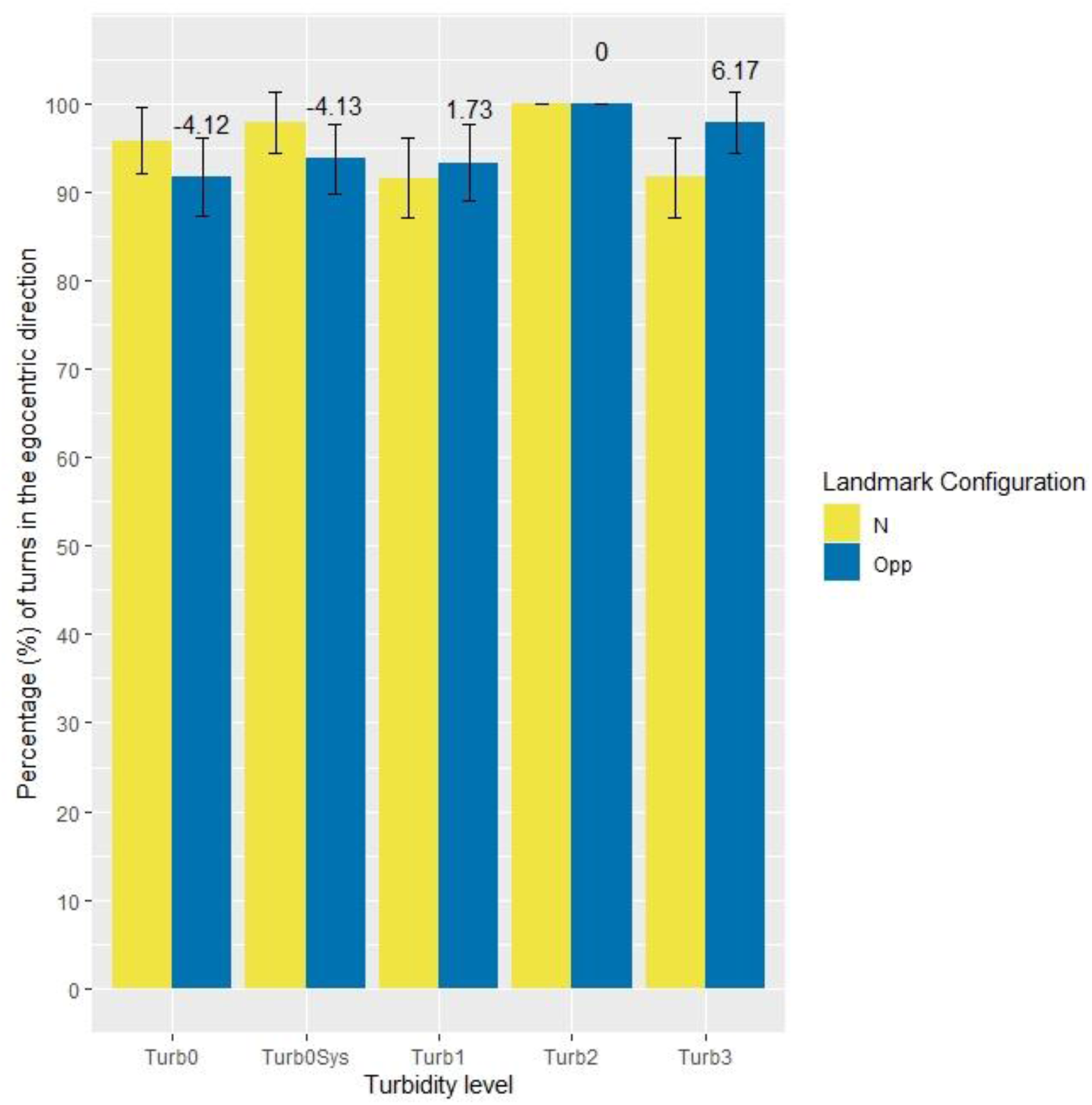
Plot of mean percentage (%) of total trials where cue conflict was present (blue) and not present (yellow) in which fish followed their learnt egocentric cue, at each turbidity level. Data from *Chrysiptera* cyanea. The plot shows the percentage (%) of total trials in which the fish took the arm corresponding to their learnt egocentric cue in N configuration trials (yellow) and Opp configuration trials (blue). Error bars indicate 95% confidence intervals about the mean. The value above each blue bar indicates the difference in percentage (%) of egocentric turns between the N configuration trials and the Opp configuration trials at that turbidity level. A negative value indicates that fish are following the landmark cue in a higher percentage (%) of trials in Opp trials than in N trials. A positive value indicates that fish are following the egocentric cue in a higher percentage (%) of trials in Opp trials than in N trials. It is crucial to understand that a negative value does *not* indicate that fish are following landmark cues more often than egocentric cues in Opp trials compared to N trials; fish would only be following landmark cues more often than egocentric cues were the mean value significantly lower than 50% at that turbidity, which is not the case in any of the turbidity levels.

**Figure 7.**
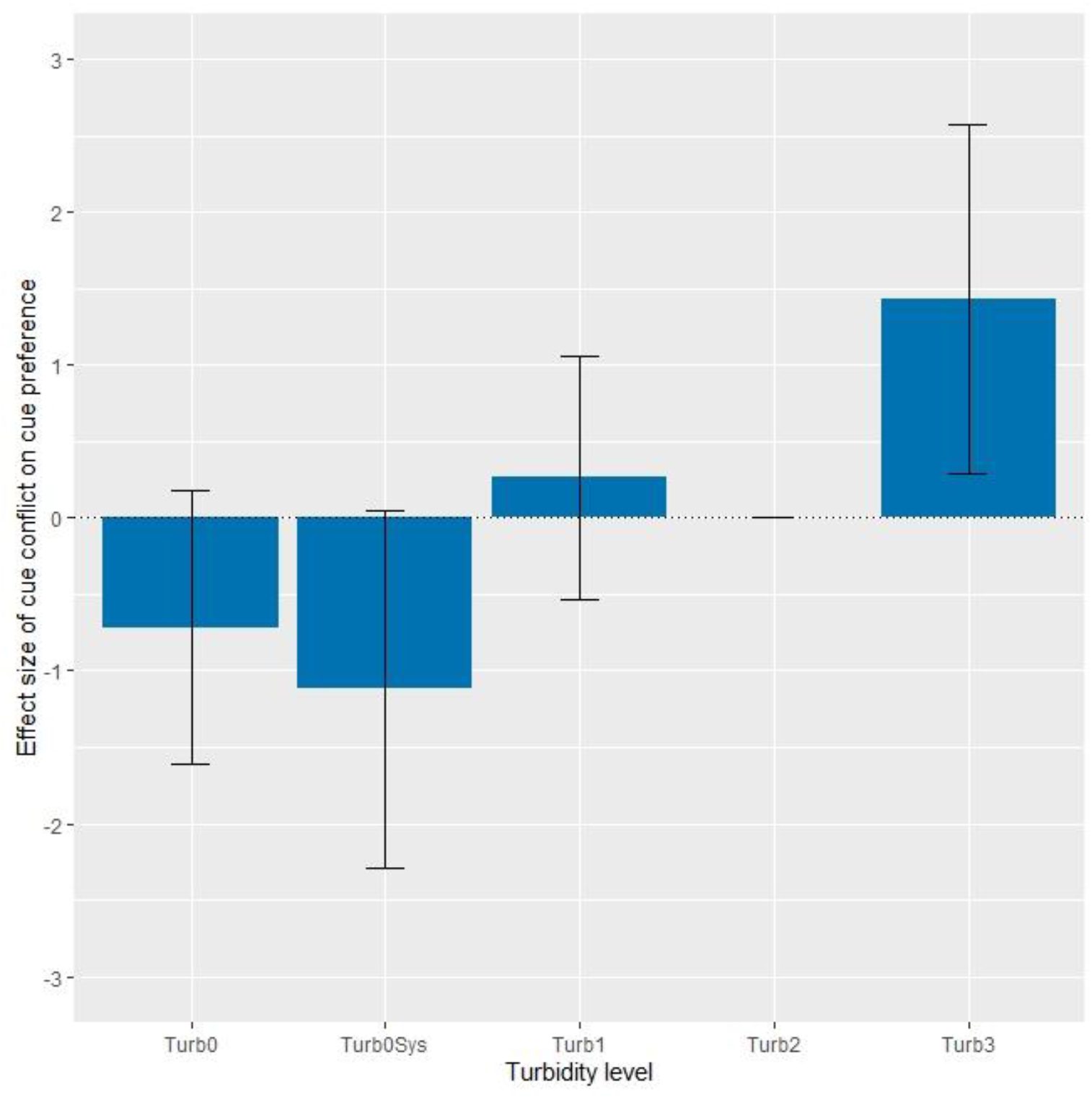
Plot of the effect of cue conflict on cue preference. Data from *Chrysiptera* cyanea. The plot shows the effect of cue conflict on the preference between egocentric and landmark cues. Error bars indicate a 95% confidence interval of the effect size. A negative effect size indicates that cue conflict (Opp configuration) causes fish to follow landmark cues more frequently than when there is no cue conflict (N configuration). A positive effect size indicates that cue conflict causes fish to follow egocentric cues more frequently than when there is no cue conflict. Negative effect sizes in Turb0 and Turb0Sys (while statistically insignificant) indicate that cue conflict in clear water causes fish to take the arm marked by the landmark more often than they would have were there no cue conflict. The positive effect sizes in Turb1 and more substantially in Turb3 (while again statistically insignificant) indicate that cue conflict causes fish to follow egocentric cues more often than they would have were there no cue conflict. The effect size is not significantly different from 0 in any of the turbidity levels (Turb0, *z* = -0.803, *p* = 0.422; Turb0Sys, *z* = -0.955, *p* = 0.340; Turb1, *z* = 0.333, *p* = 0.739; Turb2, *z* = 0.000, *p* = 1.000; Turb3, *z* = 1.258, *p* = 0.208). The pattern of these results, while statistically insignificant, suggests that in clear water cue conflict causes the preference for egocentric cues over landmark cues to weaken slightly (while still being greatly in favour of egocentric cues), while cue conflict in turbid water causes the preference for egocentric cues to strengthen. This graph is best understood when viewed in conjunction with the values above each blue bar in Figure 6, noticing the correspondence between the sign (+/-) and magnitude of those values in Figure 6 and the sign (+/-) and magnitude of the effect size of cue conflict in Figure 7.

### Trajectory analysis

Analyses of movement trajectories during testing show a clear effect of elevated turbidity on multiple movement parameters (Figure 8). LMMs were utilised for the following analyses, with the assumptions of linearity, homogeneity of variance and normality being met utilising untransformed data, unless stated otherwise. Elevated turbidity significantly reduced the average speed (*F*_*4,103*_ = 5.493, *p* < 0.001), with fish swimming more slowly in Turb2 (*t*_*174*_ = -2.509, *p* = 0.013) and Turb3 (*t*_*175*_ = -3.010, *p* = 0.003) than in Turb0 (Figure 8.a). Similarly, elevated turbidity significantly reduced the average mobile speed (*F*_*4,100*_ = 10.602, *p* <0.001), with fish swimming more slowly in Turb0Sys (*t*_*26*_ = 22.790, *p* = 0.014), Turb2 (*t*_*198*_ = -2.962, *p* = 0.003) and Turb3 (*t*_*199*_ = -4.178, *p* <0.001) than in Turb0 (Figure 8.b). Elevated turbidity significantly reduced average acceleration (*F*_*4,101*_ = 9.531, *p* < 0.001), with lower acceleration in Turb0Sys (*t*_*265*_ = -2.623, *p* = 0.009), Turb1 (*t*_*202*_ = -2.101, *p* = 0.037), Turb2 (*t*_*192*_ = -4.156, *p* < 0.001) and Turb3 (*t*_*193*_ = -4.956, *p* < 0.001) than in Turb0 (Figure 8.c). Hence, turbidity reduced the movement speed and acceleration of subjects.

**Figure 8.**
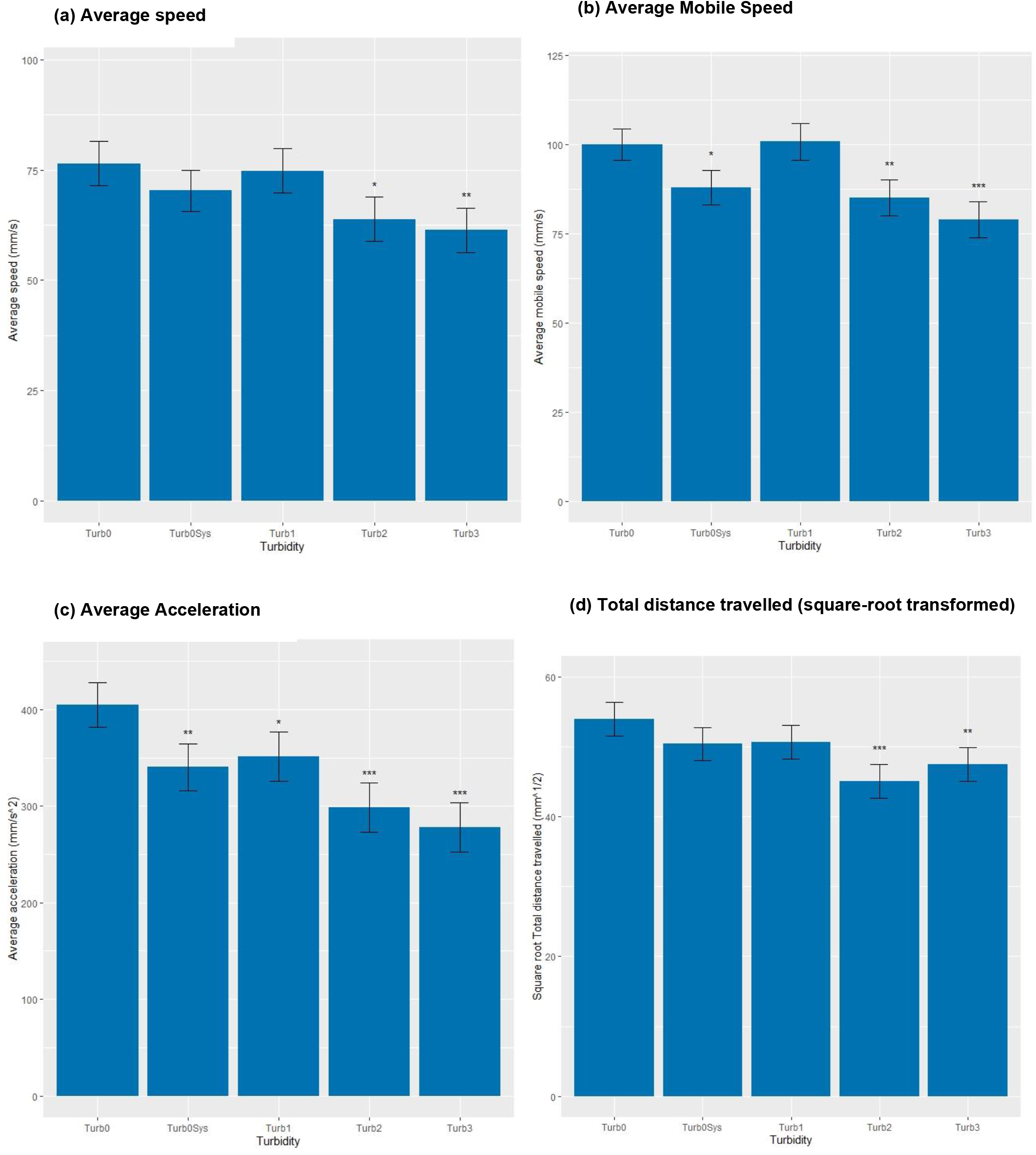
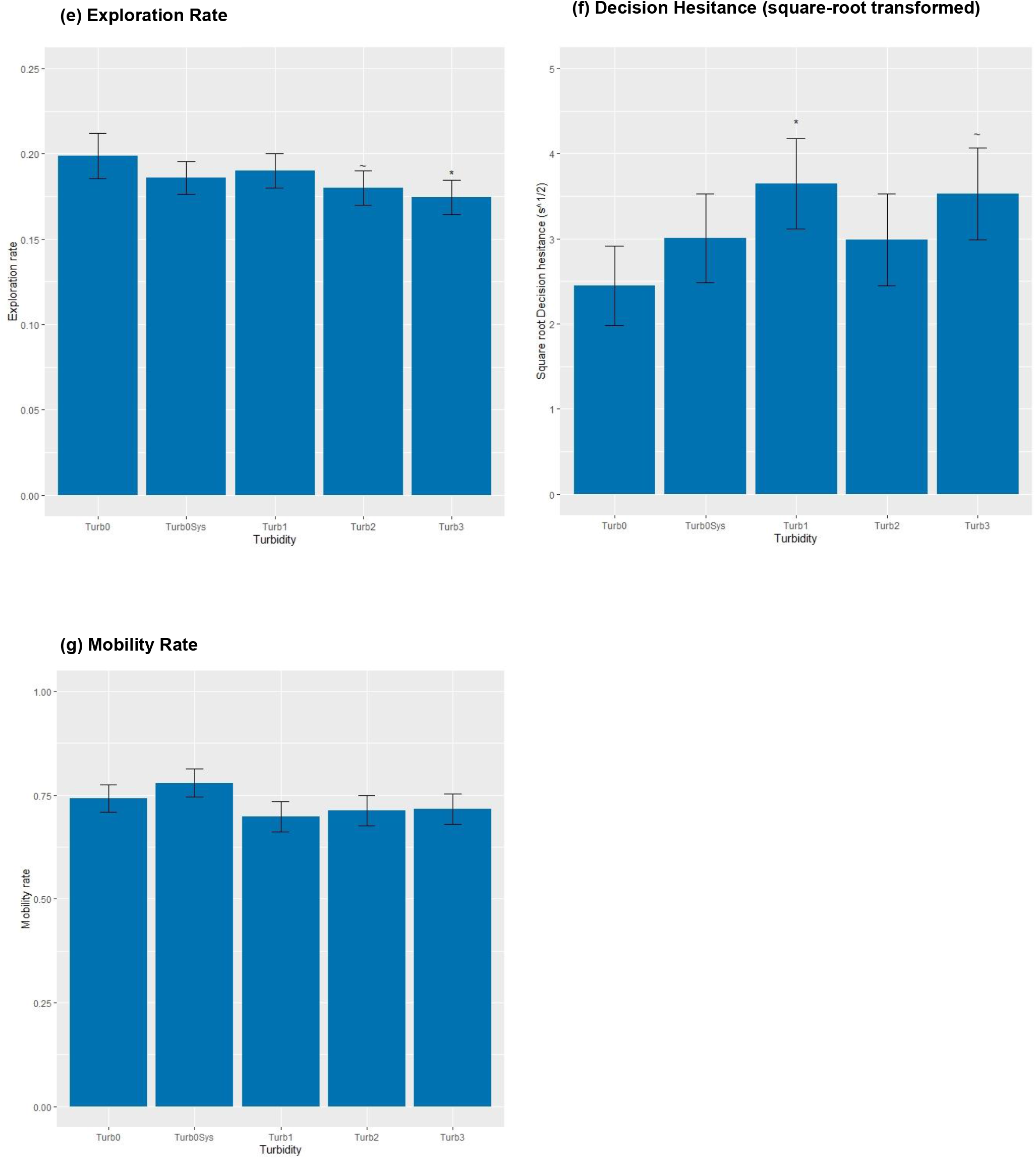
Panel displaying mean result of seven movement parameters at each turbidity level. Data from *Chrysiptera* cyanea. This panel displays the mean result for seven parameters underpinning the movement and decision behaviour of fish. Error bars indicate 95% confidence intervals. The * above a bar indicates significance with 95% confidence, ** with 99% confidence, and *** with 99.9% confidence. The ∼ above a bar indicates significance with 90% confidence. **(a)** Average speed: the mean speed (mm/s) of individuals across the entire trial, **(b)** Average mobile speed: the mean speed (mm/s) of individuals exclusively in the portions of the trial where they were not still, **(c)** Average acceleration: the mean acceleration (mm/s^2^) across the entire trial, **(d)** Square-root transformed total distance (mm^1/2^): the total distance travelled in a trial, after a square-root transformation to restore normality for analysis, **(e)** Exploration rate: the proportion of the 50×50mm squares comprising the entire arena that the fish entered at any point during the trial **(f)** Square-root transformed decision hesitance (s^1/2^): the time taken to take a turn after entering the maze, after a square-root transformation to restore normality for analysis, **(g)** Mobility rate: the proportion of the entire trial in which the fish was moving.

Fish took shorter paths at higher turbidities (*F*_*4,75*_ = 6.366, *p* < 0.001), with a square-root transformation required to normalise the response (Figure 8.d). Turbidity significantly reduced the exploration rate of fish (*F*_*4,103*_ = 3.66, *p* = 0.008), with fish exploring less of the arena in Turb3 (*t*_*193*_ = -2.446, *p* = 0.015) than in Turb0 (Figure 8.e). An analysis of space-use by Fish 3 revealed that the individual spent significantly longer at certain bicoordinates in the maze than others across all turbidities (*F*_*100,6879*_ = 10.831, *p* < 0.001), most notably within the arm which they had been trained to turn towards and at the decision junction (Figure 9). Turbidity significantly increased decision hesitance when the response was square-root transformed to partially restore normality (*F*_*4,106*_ = 2.572, *p* = 0.042) (Figure 8.f). While this transformation was the most successful in restoring normality, it was not perfect, and consequent caution should be taken when accepting this result. Following a cube transformation to restore normality, turbidity was found to have no influence on the mobility rate (*F*_*4,101*_ = 1.606, *p* = 0.179) (Figure 8.g). Thus, fish take shorter, less explorative paths in higher turbidities, with decisions potentially taking longer.

**Figure 9.**
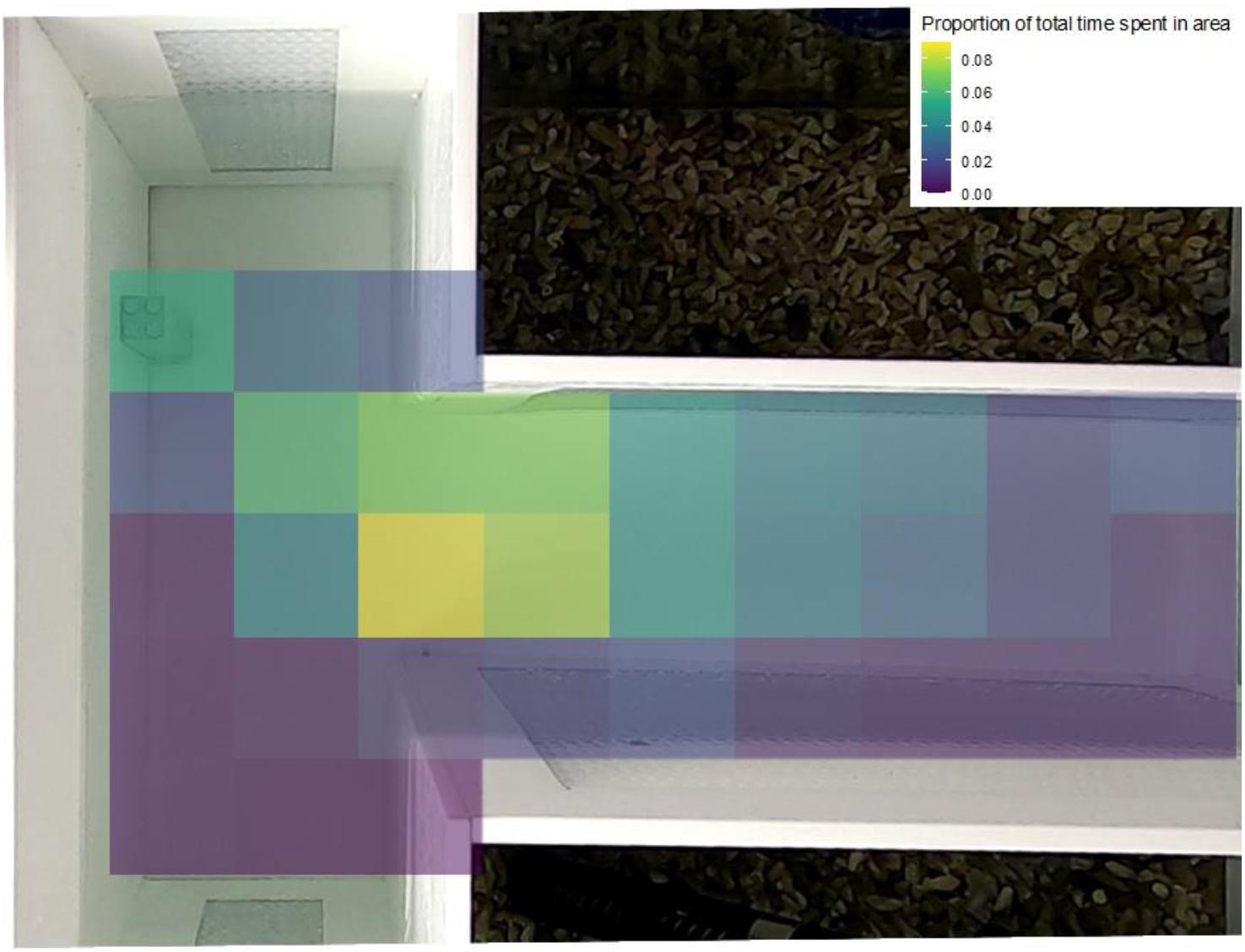
Exploration heat map for Fish 3. Data from *Chrysiptera* cyanea. This plot shows an exploration heat map for Fish 3, displaying the proportion of time spent in each 50×50mm square comprising the arena as a blue-yellow colour gradient, superimposed over an image of the T-maze in the N configuration. A yellower colour indicates a higher mean proportion of the total trial time spent in that 50×50mm square, while a bluer colour indicates a lower mean proportion of the total trial time spent in that 50×50mm square. The exploration heat map for Fish 3 displays the most yellow colours in the right arm of the T-maze, which corresponds to the direction which that fish was trained to turn in, and at the decision junction of the T-maze, namely the region just before the branching of the maze arms. A slightly yellower colour is seen on the right flank of the stem of the T-maze when compared with the left flank, hinting at a degree of lateralisation. The data shown in this plot are averaged across all turbidity conditions. This was deemed a suitable decision as turbidity was found to have no influence on the bicoordinates which fish spent the most time in (*F*_*290,6879*_ = 0.883, *p*=0.922).

Landmark configuration had no influence on any of the movement parameters analysed. None of average speed, mobile speed, acceleration, exploration, mobility rate or square-root transformed path distance were influenced by landmark configuration (respectively, *F*_*4,103*_ = 5.493, *p* = 0.902; *F*_*4,100*_ = 10.602, *p* = 0.993; *F*_*4,101*_ = 9.531, *p* = 0.980; *F*_*4,103*_ = 3.663, *p* = 0.148; *F*_*4,101*_ = 1.606, *p* = 0.515; *F*_*4,101*_ = 6.048, *p* = 0.788). Landmark configuration almost had a significant effect on square-root transformed decision time (*F*_*4,106*_ = 2.572, *p* = 0.081), indicating that conflict between egocentric and landmark cues may lead individuals to hesitate more in navigational decisions. These results suggest that individuals may have been entirely ignoring landmark cues whilst completing this task.

### Speed-efficiency trade-off

An analysis of the influence of turbidity on the range and distribution of solutions in speed-efficiency trade-off space found the effect of turbidity to be inconsequential. Following removal of outliers and a cube transformation to restore normality, a linear model reported no significant effect of turbidity on the area of the 95% confidence interval ellipses encapsulating the trade-off solutions at each turbidity (*F*_*4,20*_ = 1.696, *p* = 0.190). However, the size of these areas varied significantly between fish (*F*_*1,20*_ = 8.251, *p* = 0.009). These findings indicate that the range of solutions adopted was unaffected by turbidity, but that different fish exhibited different degrees of variability in their solutions to the trade-off. No interaction between turbidity and fish ID was reported (*F*_*4,20*_ = 1.430, *p* = 0.261), indicating this insignificant influence of turbidity was consistent across fish.

If elevated turbidity reduces the overlap area between two ellipses such that the overlap is smaller than that between two ellipses at Turb0, turbidity is inducing a shift in the solutions in speed-efficiency trade-off space. To obtain the control, constituting the overlap area between two ellipses at Turb0, the Turb0 dataset was halved and an ellipse drawn for each half of the dataset, before the overlap area between these ellipses was calculated. Ideally, these two Turb0 ellipses would be drawn from two full Turb0 datasets, but temporal and logistical constraints did not allow this. Following removal of outliers and a square-root transformation of overlap areas to restore normality, turbidity was found to have no significant effect on the size of the overlap area between ellipses (*F*_*10,44*_ = 0.495, *p* = 0.884). The size of overlap areas varied significantly between fish (*F*_*1,44*_ = 15.722, *p* < 0.001), although no interaction was reported between turbidity and fish ID (*F*_*10,44*_ = 0.675, *p* = 0.742), suggesting the insignificant effect of turbidity on the overlap area between ellipses was consistent across fish. These findings indicate that turbidity is not causing the distribution of speed-efficiency solutions to this navigational task to shift in trade-off space.

### ToxTrac Performance

To assess the viability of ToxTrac as a tracking tool in turbid conditions, the visibility of fish was compared across turbidities, first utilising data collected by ToxTrac, then utilising data collected manually from a random sample (n=120) of videos. Following a cube transformation to restore normality, turbidity was found to significantly reduce the visibility of fish in ToxTrac (*F*_*4,100*_ = 4.512, *p* = 0.002), with lower visibility in Turb3 than in Turb0 (*t*_*202*_ = -3.115, *p* = 0.002). However, following a square transformation to restore normality, turbidity was found to have no effect on the visibility of fish in the manually-collected naked-eye data (*F*_*4,110*_ = 0.228, *p* = 0.922), with no significant difference between any of the turbidity levels. These conflicting findings suggest that the reduction in visibility in Turb3 reported by ToxTrac is due to turbidity obscuring the fish from the tracking software, rather than due to confounding factors such as fish spending less time in the arena at higher turbidities. It is necessary to consider the 8% fall in visibility in Turb3 when interpreting our findings.

### Systematic turbidity increase

To elucidate a potential effect of the systematic turbidity increase, parameters were compared between Turb0 and Turb0Sys. Utilising LMMs, average speed, mobile speed, acceleration, visibility, exploration rate, distance travelled and entrance latency were all significantly lower in Turb0Sys than in Turb0 (respectively, *t*_*182*_ = -4.377, *p* < 0.001; *t*_*182*_ = -5.749, *p* < 0.001; *t*_*182*_ = -5.714, *p* < 0.001; *t*_*182*_ = -3.008, *p* = 0.003; *t*_*182*_ = -2.294, *p* = 0.023; *t*_*185*_ = -2.596, *p* = 0.010; *t*_*57*_ = -2.277, *p* = 0.027), indicating these parameters were influenced by the systematic turbidity increase. The systematic turbidity increase had no influence on mobility rate, turn latency or decision time (respectively, *t*_*182*_ = 0.648, *p* = 0.518; *t*_*180*_ = 0.534, *p* = 0.594; *t*_*179*_ = 0.992, *p* = 0.323). Hence, the systematic increase in turbidity influenced multiple parameters underpinning local navigation.

### Intrasession variation

In the experimental phase, fish had to complete four consecutive unrewarded trials before any reward was provided. Consequently, one might expect motivation to fall over the course of a session. Utilising LMMs, a reduction in average speed (*F*_*1,469*_ = 14.327, *p* < 0.001), mobile speed (*F*_*1,469*_ = 9.010, *p* = 0.003) and acceleration (*F*_*1,469*_ = 30.063, *p* < 0.001), and an increase in decision hesitance (*F*_*1,460*_ = 4.768, *p* = 0.030) were reported over the course of a session. No changes were seen in mobility rate (*F*_*1,469*_ = 1.322, *p* = 0.250), exploration rate (*F*_*1,469*_ = 0.099, *p* = 0.920), distance travelled (*F*_*1,469*_ = 0.574, *p* = 0.449), entrance latency (*F*_*1,468*_ = 0.155, *p* = 0.694) or turn latency (*F*_*1,460*_ = 2.839, *p* = 0.093). Consequently, a fall in motivation over the course of a session may have occurred.

## DISCUSSION

This study provides the first insight into the effect of turbidity in shaping the preferences of reef fishes between different navigational cues. The analysis suggests that turbidity has no significant influence on the preference between visual and egocentric information in a short-range navigational task. Instead, subjects exhibited a stark preference for egocentric cues which was robust to changes in turbidity, committing to the arm signalled by their learnt egocentric cue in at least 90% of trials when cue conflict was present. Such extreme preference for egocentric cues was unexpected for two reasons. Firstly, coral reefs are visually rich ecosystems, governed by a multitude of visually-guided behaviours (Marshall et al. 2019; Kamermans & Hawryshyn 2011). Consequently, one would expect visual information to hold substantial weight when informing navigational decisions. Secondly, the reliability of information supplied by egocentric cues was equal to that provided by landmarks in training, with both cues being associated with equal strength to a food reward.

The polarity in favour of egocentric cues begs the question of whether the landmark was ignored altogether. Of the six movement parameters, none were influenced by landmark configuration. This apparent inattention to landmark cues is striking. Such indifference to landmark cues may have multiple sources. Firstly, while the two cues had identical correlations with food provision in training, the landmark cue may have become perceived as unreliable in the experimental phase, during which the landmark was frequently moved between arms. The mobility of the landmark may have reduced its perceived reliability as an indicator of food provision (Odling-Smee & Braithwaite 2003b). Secondly, a strong egocentric preference is often characteristic of overtraining (Kealy et al. 2008), although this is unlikely given the maximum number of training sessions any one fish experienced prior to testing was five. Thirdly, one might suggest that this egocentric preference may have not been egocentric at all, but rather visual, with fish identifying a landmark in the testing room and learning the location of food provision relative to that landmark. However, the softbox lights placed in front of each tank blocked access to visual features beyond the tank, rendering this possibility unlikely. One exception was the experimenter’s face, which was visible to fish in front of the lights. Should fish have identified a facial feature with a consistent position relative to the training arm, fish could have been orienting relative to this landmark. Capacity for human facial recognition has indeed been demonstrated in teleosts (Newport et al. 2016; Newport et al. 2018).

With attendance to visual landmarks being inapparent from the trajectory analysis, the effect of landmark configuration may be more subtle than initially hypothesised. While there was no significant difference in the percentage of turns in the egocentric direction between N trials and Opp trials at any turbidity level, these differences, insignificant as they were, did display the hypothesised pattern. In clear water, a higher frequency of landmark turns was seen in Opp trials than in N trials. Conversely, at higher turbidities (with the exception of Turb2, where every turn was in the egocentric direction), a higher frequency of egocentric turns was seen in Opp trials than in N trials. These data suggest that a shift in cue preference may have been occurring in response to rising turbidity on a far finer scale than originally hypothesised. This shift in cue preference was hypothesised to operate between two polarities, whereby high quality visual information in clear water would pull fish to a strong landmark preference (∼20% egocentric turns), and poor quality visual information in turbid water would push fish to a strong egocentric preference (∼80% egocentric turns). Thus, instead of cue preference flipping from one extreme to the other as turbidity rises, this shift occurred on a subtler scale, within the confines of a strong egocentric preference (90% egocentric turns) and an even stronger egocentric preference (100% egocentric turns). With this pattern existing on a small (albeit statistically insignificant) scale in response to the range of turbidities employed here, future work exploring this modification of navigational cue preference over a broader range of turbidities would be enlightening in the face of rising coastal turbidity.

Results from the trajectory analysis suggest a substantial role for turbidity in influencing the movement behaviour of reef fishes. Fish swam slower at higher turbidities, taking longer to make navigational decisions. This combination of slower movement and decision speed may increase exposure to predators if such decisions occur in exposed areas of the reef. With such changes being detectable even within the limited area of the T-maze, the implications of these changes may be considerable in the broader ecological range that wild fish operate in. While anecdotal, the exploration heat map produced for Fish 3 may suggest that these navigational decisions are deliberated at the decision junction of the T-maze, reflected by the highest proportion of time being spent in the bicoordinates at the branching of the T-maze; but this will remain conjecture until further testing.

The apparent increase in decision hesitance does not come at the cost of efficiency, with fish taking shorter, less explorative paths in more turbid water. Such an increase in path efficiency may begin to counter the potential increased predator exposure arising from increased hesitance. This increase in navigational efficiency has interesting implications for the necessity (or lack thereof) of integrating information from multiple cues for efficient navigation. In clear water, fish had access to both visual and egocentric information. In turbid water, the quality of visual information was lessened. Therefore, the fact that fish not only maintained but increased their navigational efficiency in spite of the degradation of visual information implies a redundant role of visual landmarks in guiding navigational efficiency.However, the ecological applicability of this conclusion is limited. Given the short distances that fish were confined to operate within while in the T-maze, it is unsurprising that egocentric information alone provides sufficient guidance in this simple task. One should expect the degradation of visual information to only impart substantial costs to navigational efficiency when fish are operating over larger, more ecologically relevant distances.

Despite the mean decision speed decreasing and the mean decision efficiency increasing with turbidity, no shift in speed-efficiency trade-off space was observed. Consequently, while the rise in turbidity employed in this study was enough to induce shifts in the aforementioned means, these changes are unlikely to be coupled with a shift in the entire distribution of solutions in trade-off space until fish are exposed to more extreme turbidities than those utilised here. Furthermore, with the range of solutions to the speed-efficiency trade-off being unaffected by a rise in turbidity, the diversity of solutions to navigational speed-efficiency trade-offs is unlikely to be eroded by visual pollution of a similar magnitude to that generated here.

While the findings presented may have implications for cue preference, movement behaviour and decision-making, there are limitations to consider when gauging the applicability of these results. Firstly, and most pertinently, logistical constraints of the experimental system demanded a systematic increase in turbidity be used, rather than the preferable randomisation of turbidity from trial to trial. With differences in multiple movement parameters between Turb0 and Turb0Sys, it is evident that this systematic turbidity increase influenced fish behaviour. Given these changes persisted after the return to clear water, future work exploring how long such modifications persist after returning to clear water would shed light on the long-term implications of periodic turbidity exposure. Furthermore, with reductions in movement and decision speeds over the course of an experimental session, motivation likely contributed noise to our analyses. Thirdly, while ToxTrac operated at a high standard in the majority of trials, this standard slipped at higher turbidities. Despite this, the findings presented would not change upon accounting for noise introduced by the 8% fall in ToxTrac detection rate in Turb3, because statistically significant differences between Turb0 and Turb3 were already evident in all movement parameters influenced by turbidity even before accounting for this noise. In minor exception to this, the difference in decision hesitance between Turb0 and Turb3 would likely become statistically significant upon accounting for the aforementioned noise.

While this study is the first to explore the influence of reduced visual range on preference between visual and egocentric cues for navigation, turbidity has influences beyond limiting the perceptual range. The differential alteration of the spectral environment by different sources of turbidity likely has profound effects on the attendance of fish to landmarks of different colours. With non-algal particulates (NAP) attenuating blue wavelengths, landmarks which would have otherwise appeared blue will appear less salient during NAP plumes. Similarly, with CDOM attenuating strongly in the blue-ultraviolet range, ultraviolet signals may become imperceptible in CDOM plumes. Consequently, one might expect fish to attend differently to landmarks of different colour depending on what material is generating the local turbidity. An exploration of this may be conducted through employing the experimental design described herein utilising a range of landmark colours and turbidity sources, and is of paramount importance if we are to understand how landmark use in navigation is to be influenced in the coming decades of increasing anthropogenic visual pollution.

## CONCLUSIONS

This study reports no significant effect of turbidity on the preference of damselfish between different navigational cues. Instead, fish exhibited a striking preference for egocentric cues which was robust to changes in turbidity. The hypothesised shift in cue preference may be occurring on a subtler scale than originally thought. Turbidity has profound implications for the movement behaviour of damselfish, with these implications likely being of substantial magnitude on the ecological scale of a coral reef.

## ACKNOWLEDGEMENTS

We thank the Wytham Field Station Fish Group for their insight and guidance. We thank Christine Soper for help with animal husbandry and John Hogg for the construction of the T-maze. This research was funded by the Human Frontier Science Program grant number RGP0016/2019. Cait Newport was funded by the Leverhulme Trust Early Career Fellowship.

## SUPPORTING INFORMATION

All Supporting Information may be found online at **XXXXX**.

